# Evo 2 as a classification machine: evidence from in-context learning and mechanistic interpretability

**DOI:** 10.64898/2026.09.30.755330

**Authors:** Lorenzo Bertolini Agnoletto, Fabiola Curion, Mauro Petrillo, Gabriele Leoni, Michele Ronco, Victoria Ruiz Serra, Sergio Consoli, Mario Ceresa

## Abstract

In-context learning (iCL) is an emergent capability of Large Language Models (LLMs), allowing them to perform new tasks at inference time using prompt-injected examples. While extensively studied in LLMs, the boundaries of its capabilities and underlying mechanisms remain poorly characterised in genomic Language Models (gLMs). Here, we map the operating regime of Evo 2, a nucleotide-level foundation gLM, across five binary classification tasks spanning biological and artificial sequences. We observe robust iCL on shorter natural sequences (F1=0.902 for miRNA, 0.785 for Toxins), degrading with sequence length, and collapsing at kilobase scale. Strikingly, we find no benefit in model scaling, as the 7B model systematically outperforms the 40B variant. We further find that perplexity, a widely used gLM performance proxy, poorly predicts accuracy. Mechanistic interpretability reconciles these observations: the logit-lens profiling suggests a prediction-generalisation trade-off, while Jacobian Scope indicates models might track prompts’ structure rather than the signal-carrying content.

---

Genomic foundation models (gLMs) pretrained autoregressively on nucleotide sequences have recently emerged as a powerful paradigm for decoding the genomic language [1-4]. Architectures such as HyenaDNA [5], Evo [6] and GROVER [7] have demonstrated that large-scale nucleotide pretraining encodes rich sequence-level representations transferable to diverse downstream tasks, ranging from genomic classification to zero-shot variant prediction and generative sequence design. Evo 2 [3, 8] expanded this frontier using the computationally efficient StripedHyena 2 architecture trained on over 9 trillion multispecies tokens, offering a large suite of models, with up to 1M-token context windows.

Conventionally, to evaluate whether such models have successfully assimilated biological sequence structure, the field has relied on perplexity (Methods) [3, 9, 10], a measure of how well a language model predicts the next token in a sequence. Lower perplexity indicates that the model is less ‘‘surprised’’ by the test sequence [3]. To adapt gLMs to downstream tasks, researchers generally relied on likelihood-based scoring. The approach typically compares distributions of scores across sets of sequences, making it difficult to classify a single sequence without resorting to a supervised classifier trained on extracted embeddings [3]. These approaches are poorly suited for single-sequence classifications, especially in the absence of large quantities of labelled data.

To alleviate this issue, researchers recently started adopting gLM-suited in-context learning (iCL) solutions [11-14]. iCL is an emerging ability observed in large language models (LLMs) [15, 16], allowing them to perform novel tasks from annotated contextual examples injected in the prompt, alongside a query, without the need for weight updates [15, 16]. A recent work showed that Evo 2 models can function as in-context learners [11] by comparing the perplexity of candidate label completions appended to the query. However, this initial characterisation focused almost exclusively on synthetic, ‘‘puzzle-like’’ sequence tasks [11]. Consequently, the extent to which genomic iCL generalises to *real-world* biological classification, how it scales across model parameter sizes, and how it is influenced by sequence length remain unaddressed, together with the underlying inner mechanisms of the model.

Here we map this operating regime through a structured test set, with a significantly stronger focus on natural and biologically plausible sequences that span a gradient of biological complexity, evolutionary dynamics, and real-world relevance, namely microRNAs (miRNAs), toxin-encoding genes, and antimicrobial resistance (AMR) genes. We complement the real-world tasks with a synthetic classification task based on TATA-box promoter regions at two length regimes (20-100 bp and 1-3 kb). We evaluate two Evo 2 models alongside two state-of-the-art medical LLMs, systematically varying the number of in-context examples per class (*N* = 1 to 256). Lastly, to aid interpretation of the observed model behaviour, we conduct mechanistic interpretability analyses — logit lens and Jacobian Scope — to ask not only *whether* iCL succeeds but *why* it succeeds or fails.

Our contributions are fourfold. *(i)* We provide the first broad, multi-task, multi-scale characterisation of genomic iCL on real biological classification problems, moving beyond the synthetic, puzzle-like tasks that dominated prior work. *(ii)* Using a matched synthetic control that holds the discriminative signal fixed while varying only sequence length, we show that sequence length — not biological complexity — is what breaks perplexity-based scoring under a fixed compute budget, giving practitioners a clear operating regime. *(iii)* Furthermore, we show that perplexity, the field’s default performance proxy, is a weak predictor of iCL accuracy, and can become *anti-correlated* with it as the sequence length increases. *(iv)* Finally, we show that scaling from 7B to 40B parameters degrades rather than improves iCL, and offer a mechanistic account in which larger models commit to next-token prediction too early in depth and, on failing tasks, track the prompt’s positional scaffold rather than its biological signal.

## 1. Results

Classification followed a previously proposed perplexity-scoring protocol [11] (Methods, Figure 1), where each DNA sequence is paired with a buffer and a class-label run, each spanning 24bp nucleotides. A model then classifies by scoring the candidate label completions appended to the in-context prompt and selecting the lower-perplexity class, requiring no weight updates.

**Figure 1.**
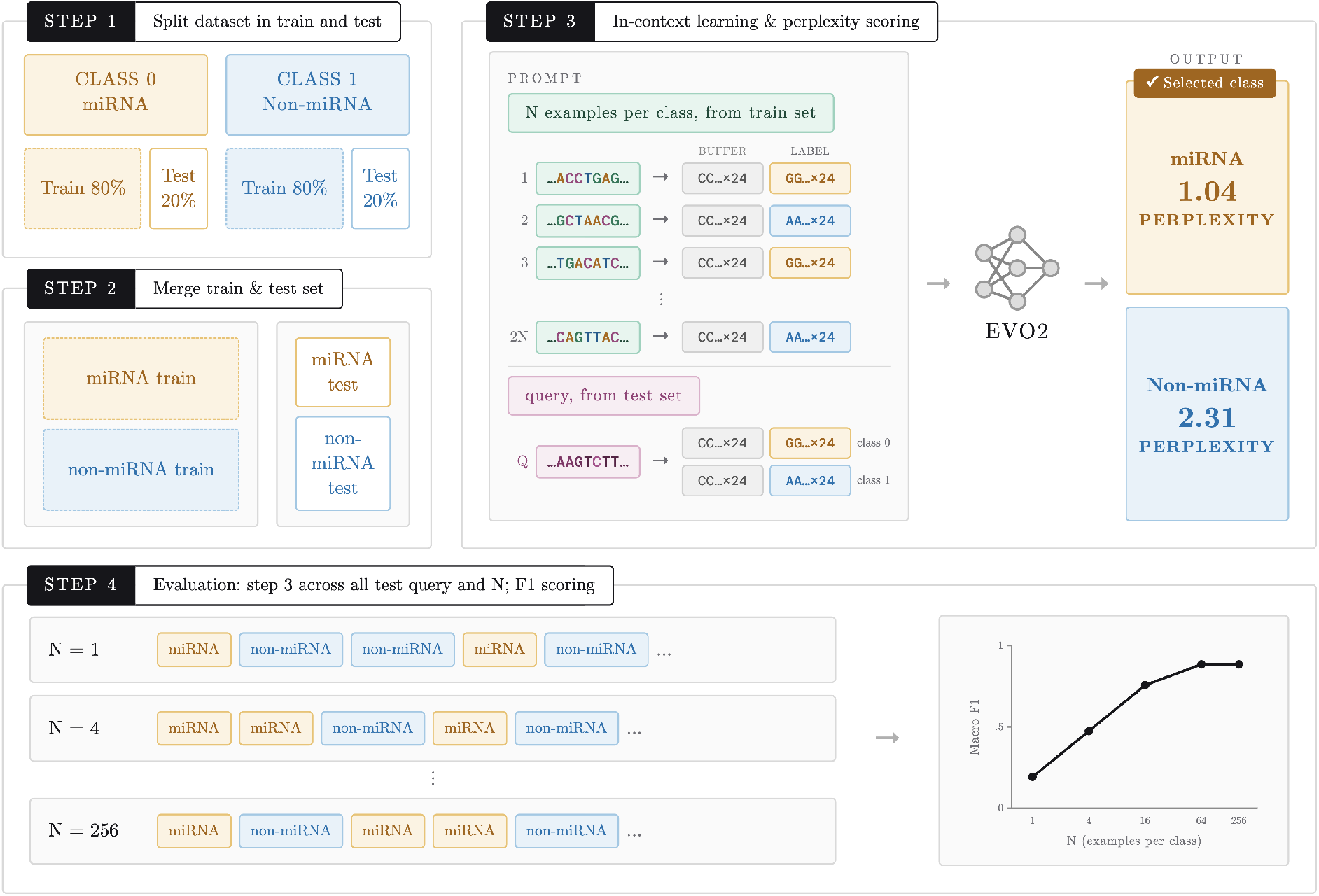
Experiment structure. Schematic representation of the main steps composing the experimental structure, exemplified with a single Evo 2 model and microRNA sequences. **Step 1**, each class pool (class 0, miRNA; class 1, non-miRNA) is split independently into train and test sets. **Step 2**, per-class splits are merged into balanced train and test sets. **Step 3**, *N* examples per class from the train set are concatenated with a test set query, each encoded as a [sequence] *→* [buffer_run] [label_run] pair. Frozen Evo 2 scores both candidate labels, and the lower-perplexity candidate is shown as the predicted class. **Step 4**, step 3 is repeated for each test query, with different values of *N*; predictions are then evaluated against ground truth with Macro F1.

Experiments were conducted on two Evo 2 models with a 1M context window — 7B, 40B — alongside two state-of-the-art instruction-tuned medical large language models, namely MedGemma 1.5 4B and MedGemma 27B text-only. We evaluated each model across five binary classification tasks grouped into two tracks: a biological track (made of three sequence classes, i.e., miRNA, toxin-encoding sequences, and AMR genes) and a synthetic control track (made of one sequence class, i.e., TATA-box sequences at 20-100 bp and 1-3 kb). A summary of the sequence classes is provided in Table 1.

**Table 1.** Overview of the natural and artificial sequence classes used to evaluate Evo 2’s in-context learning capabilities.

| Dataset | Type | Sequence Characteristics | Selective / Evol. Pressure | Primary Rationale | Translational Relevance |
| --- | --- | --- | --- | --- | --- |
| microRNAs | Natural | Short (~22 nt mature; ~60-110 nt precursor); structurally constrained hairpin | Moderate-to-high conservation; structural constraint dominates | Tests generalisation from sequence to structural/regulatory grammar | Biomarkers/therapeutic targets in disease; basic regulatory biology |
| Toxin-encoding genes | Natural | Diverse virulence factor families; variable length/structure | Variable; often mobile/plasmid-associated | Tests recognition (not generation) capability relevant to biosecurity | Underlies sequence-based biosecurity screening frameworks |
| AMR genes | Natural | Functionally diverse (β-lactamases, efflux pumps, target-modifying enzymes) | Rapid evolutionary turnover; frequent horizontal gene transfer | Tests generalisation across a functionally-defined but sequence-diverse class | Genomic AMR surveillance; WHO-recognised global public health threat |
| TATA-box promoters | Artificial | Short, strongly conserved consensus motif; well-defined positional grammar | Strong purifying selection; canonical, near-invariant motif | *Controlled benchmark:* tests whether iCL can learn a known, explicit motif under near-ideal conditions | Interpretability anchor; baseline for isolating raw pattern-recognition capability |

### 1.1. iCL performance on short natural sequences

On short natural sequences, represented here by miRNA and toxin classification datasets, Evo 2 7B achieved the highest overall performance across all tested models (Table 2). For miRNA classification, Evo 2 7B reached a macro F1 score of 0.902, outperforming Evo 2 40B. This advantage was even more pronounced in toxin classification, where Evo 2 7B achieved an F1 of 0.785 at high *N*, substantially outperforming Evo 2 40B (F1 = 0.645).

**Table 2.** Peak macro F1 across models and tasks. Values represent the maximum macro F1 achieved by each model (rows) in each dataset (columns) across the evaluated range of *N* (1-256 per class). Cell colour coding reflects the F1 score (warmer/red = higher, cooler/blue = lower), standardised across all results in the table.

| Model | Natural Sequences |  |  | Artificial Sequences |  |
| --- | --- | --- | --- | --- | --- |
|  | miRNA | Toxins | AMR | TATA 20-100 | TATA 1k-3k |
| Evo 2 7B | 0.902 | 0.785 | 0.559 | 0.629 | 0.515 |
| Evo 2 40B | 0.846 | 0.645 | 0.512 | 0.564 | 0.512 |
| MedGemma 1.5 4B | 0.634 | 0.527 | 0.513 | 0.532 | 0.508 |
| MedGemma txt 27B | 0.548 | 0.524 | 0.513 | 0.534 | 0.505 |

Across both tasks, both MedGemma models performed remarkably worse than the genomic models, with F1 scores ranging from 0.524 to 0.634. These results demonstrate two key properties of the short-sequence regime: first, nucleotide-level pretraining is indispensable for effective pattern extraction, as high-level medical instruction tuning fails to compensate; second, increasing parameter scale within the Evo 2 family does not monotonically improve in-context learning performance on short sequences.

### 1.2. Shot-Scaling Dynamics and Length-Dependent Limits

To characterise how performance scales with iCL, we evaluated shot-scaling trajectories (F1 score as a function of example count *N*) across task domains and sequence lengths (Fig. 2). These experiments revealed three distinct scaling regimes. First, on short natural sequences, increasing *N* substantially improves performance, though optimal trajectories depend on both task and model scale. For miRNA, Evo 2 7B exhibited monotonic gains — climbing from F1 *≈* 0.55 at *N* = 1 to F1 = 0.90 at *N* = 256 — whereas Evo 2 40B plateaued and degraded past *N* = 64. In contrast, on the longer AMR gene sequences (~500-3,000 bp), only Evo 2 7B showed a marginal yet steady learning (reaching F1 *≈* 0.56), while all other models stagnated near chance (F1 *≈* 0.51). Second, the synthetic TATA-box control experiments demonstrated that sequence length rather than sequence complexity — imposes a critical barrier on iCL. On short sequences (20-100 bp; TATA-short), Evo 2 7B displayed a steep, effective learning curve (reaching F1 = 0.629). However, when sequence length was extended to 1-3 kb (TATA-long), performance across all models collapsed to near chance (F1 *≤* 0.515) and degraded further as *N* increased.

**Figure 2.**
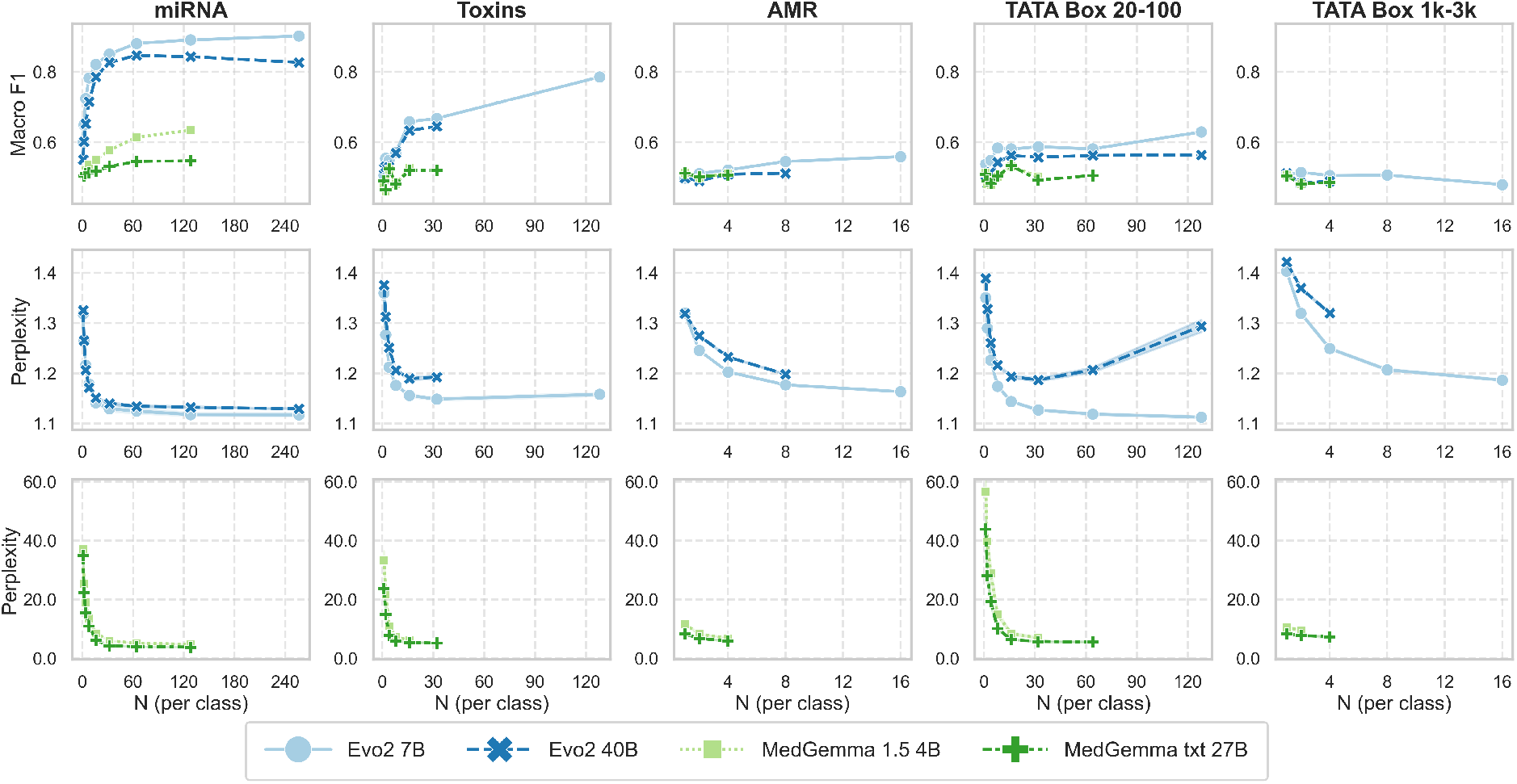
Shot-scaling of macro F1 and perplexity across models and tasks. Macro F1 (top row) and sequence perplexity (bottom row) as a function of the number of in-context examples per class *N*. Columns correspond to classification datasets and coloured lines to the evaluated models, as indicated in the legend. The top and bottom panels in each column share the same *x*-axis (*N*).

These scaling curves reflect performance within fixed compute and architectural bounds rather than fundamental saturation limits. Longer sequences drastically reduce the maximum example count *N* that can fit within a given compute budget, truncating the AMR scaling curve before a true performance ceiling could be reached. Furthermore, we ran into recursive implementation instabilities, such as non-monotonic trajectories in MedGemma, and FFT convolution errors in Evo 2. Consequently, we could not evaluate a subset of sequences since their lengths were incompatible with the FFT-based long-convolution operator underlying StripedHyena. In our own testing, we repeatedly observed numerical instability when running sequences with lengths that did not factorise favourably (e.g., were not powers of 2). A standard workaround is to pad sequences to a compatible length, but padding alters the effective input and can shift perplexity estimates in ways that would confound comparisons across instances. To preserve comparability of perplexity scores across the reported results, we therefore excluded trials presenting at least one FFT error, rather than pad multiple sequences. These observations further highlight that real-world iCL capacity is constrained as much by system-level execution limits as by intrinsic model capability.

### 1.3. Perplexity-Performance Dissociation

In gLMs, sequence perplexity — a measure of how well a model predicts subsequent tokens — is frequently reported as a proxy for downstream performance and risk assessment [3, 10]. We evaluated whether overall sequence perplexity reliably predicts in-context classification accuracy (F1 score) across varying example counts *N*. Across all tested tasks, sequence perplexity decreased roughly monotonically with *N*. However, this reduction in perplexity occurred largely independently of downstream classification success (Fig. 2). This dissociation manifested in three distinct regimes across our benchmark. For miRNA classification, Evo 2 7B and 40B exhibited co-monotonic declines in perplexity alongside F1 gains up to *N ≈* 64; past this threshold, however, Evo 2 40B perplexity continued to decrease even as F1 performance stagnated. On toxins and AMR gene classification, non-genomic LLMs exhibited steep drops in perplexity as *N* grew, despite remaining near chance performance (F1 *≈* 0.51). The long synthetic control (TATA 1k-3k) provided the most striking failure mode: while perplexity steadily decreased for all models with increasing *N*, classification F1 simultaneously collapsed.

These results strongly suggest that sequence perplexity might reflect the model’s increasing familiarity with prompt syntax and background sequence statistics, rather than its acquisition of the discriminative classification rule. Consequently, this undermines the possibility of framing raw perplexity as a surrogate for directly measuring in-context performance in Evo 2 downstream classification tasks.

### 1.4. Mechanistic insights into Genomic In-Context Processing

To uncover the internal mechanisms governing in-context learning, we applied two complementary mechanistic interpretability frameworks across the Evo 2 model suite (1B base, 7B, and 40B): logit lens analysis [17] and Jacobian influence scoring [18].

Tracking residual stream projections across network depth (at each model layer) revealed distinct representational dynamics between model scales (Fig. 3). In Evo 2 7B and the 1B base model, confident top-1 token predictions emerged mostly in the final 2-3 layers, with a mild yet consistent contribution distributed across middle layers in the 7B models, likely reflecting a gradual refinement of representations throughout the network. In contrast, Evo 2 40B established high prediction confidence prematurely, maintaining saturated top-1 predictions across early layers and the entire second half of the network. While evaluated on a representative AMR sequence, these layer-wise trajectories were consistent across multiple sequence samples and align with previously reported dynamics in the 20B spliced variant of Evo 2 [8] (Data on other sequences not shown).

**Figure 3.**
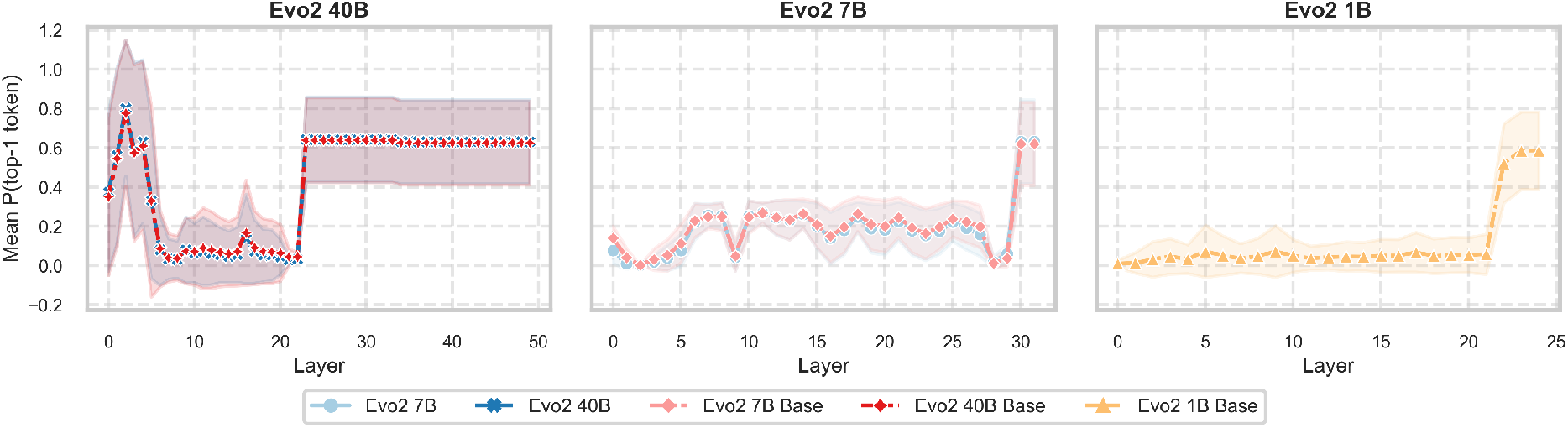
Logit-lens analysis across Evo 2 model scales. Layer-wise top-1 token probability (*P* (top-1)), averaged over input positions of a representative AMR sequence, for different Evo 2 models. The *x*-axis is model layers, *y*-axis refer to average *P* (top-1) activations, and shades represent standard deviation.

Using the Semantic Scope framework (see Methods), we quantified the causal contribution of individual prompt tokens to final label predictions. On miRNA sequence classification (Fig. 4), token influence patterns diverged sharply by parameter scale: while the 1B base model exhibited a strong recency bias (attending primarily to immediate prefix tokens), both the 7B and 40B models shifted causal weight toward early in-context prompt tokens. Crucially, across all models, causal influence failed to localise around functionally relevant biological motifs — such as uracil-rich regions characteristic of miRNAs.

**Figure 4.**
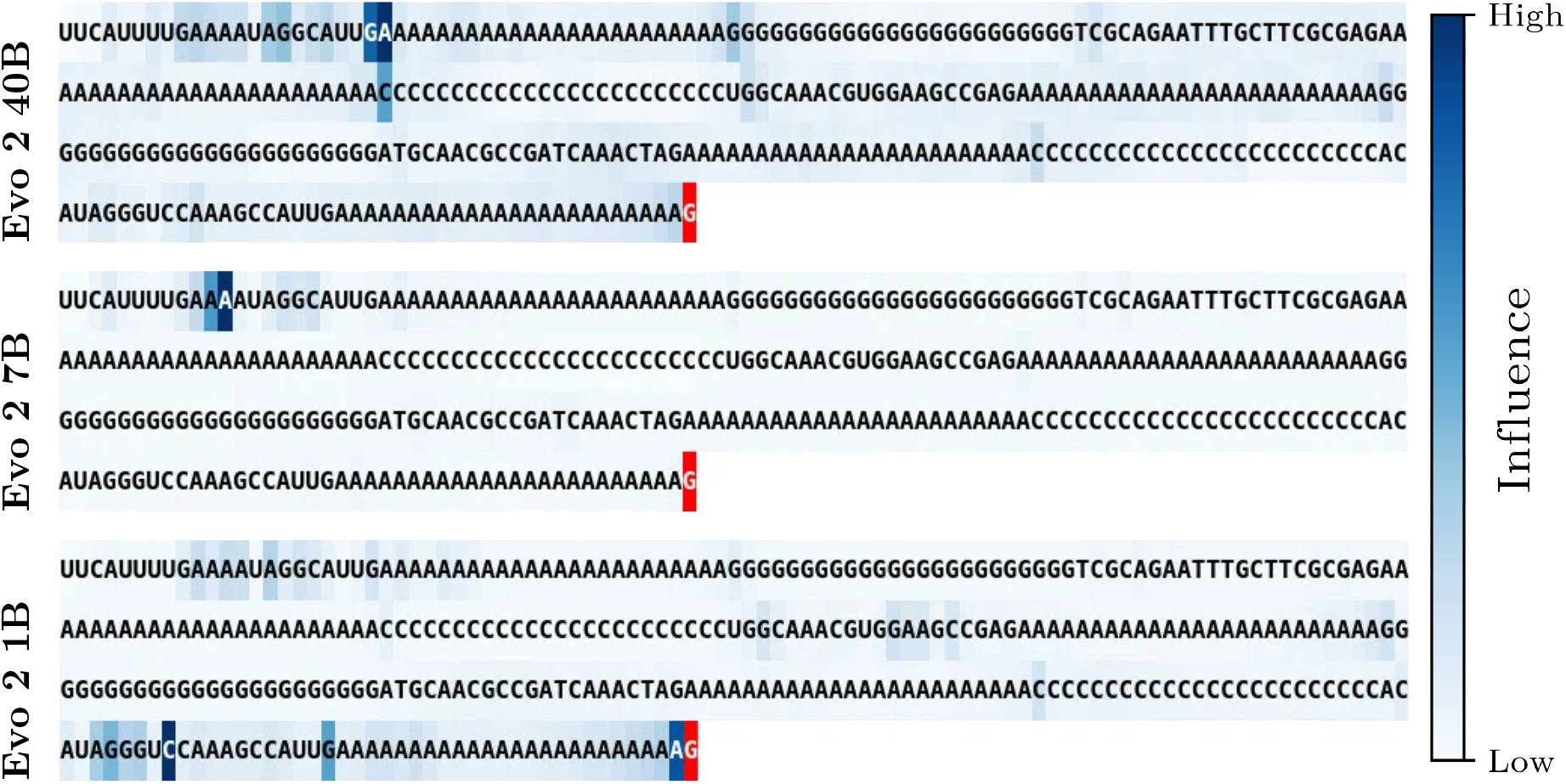
Token-level causal-influence profiles for a miRNA in-context trial across Evo 2 scales. Each row corresponds to one model (top to bottom: Evo 2 40B, 7B and 1B base). In each row, the target token (red) is the label-run token whose prediction is being explained, and the remaining tokens are the in-context prompt (context) tokens on which its influence is computed. Colour intensity encodes the Semantic Scope influence value (see Methods): how strongly each prompt token contributed to generating the target token, from low (light) to high (dark), as indicated by the shared colour bar on the right.

We then evaluated synthetic TATA-box sequences where the ground-truth signal location is known *a priori*. We computed Semantic Scope profiles across all 24 tokens encoding the label to track attribution dynamics across the completion sequence (Fig. 5). The resulting influence profiles revealed that causal contribution localised to the embedded TATA motif is minimal and virtually indistinguishable from surrounding background sequences. Instead, the dominant influence peaks recur periodically at the structural boundaries of the iCL prompt (the transitions into the label-run regions), tracking the syntactic [sequence] *→* [label_run] template rather than its semantic content.

**Figure 5.**
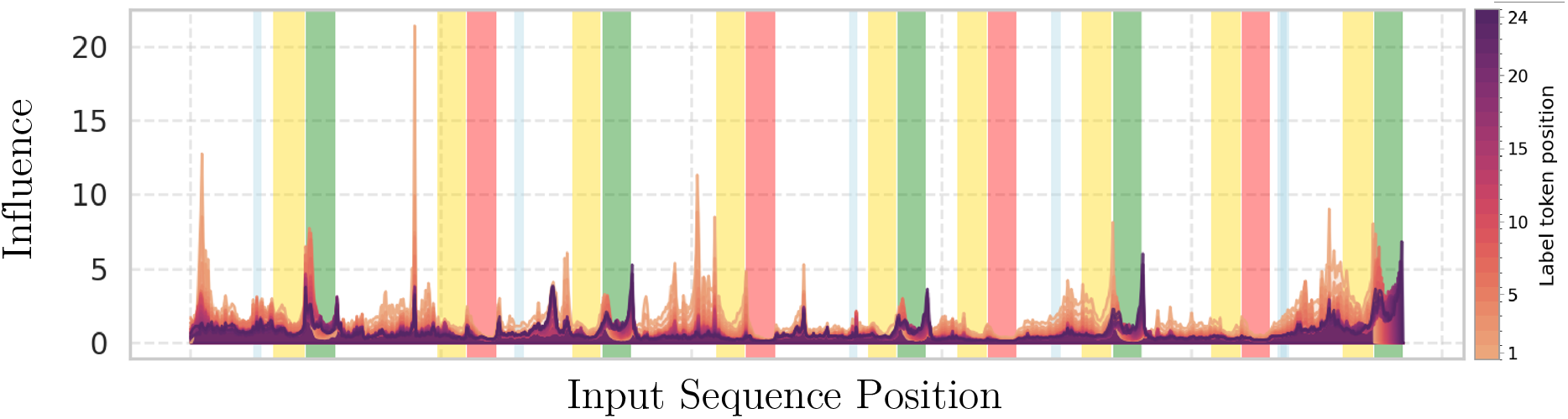
Token-level causal-influence profile for a TATA-box in-context trial (*N* = 4, Evo 2 7B). Coloured vertical bands mark the recurring regions of the prompt scaffold: blue, the embedded TATA-box motif (the only class-discriminative region); yellow, buffer (spacer) sequence; red, neutral (non-TATA) label region; green, target (TATA) label region. The overlaid curves are the Semantic Scope influence profile (see Methods) as a function of label token position; curve height is the per-token influence read off the *y*-axis, and the band colours are categorical region labels rather than a magnitude scale.

By design, we know *where* models should focus attention to solve the task: the TATA-box sequence. Contrary to the miRNA investigation, we gathered Semantic Scope profiles for all 24 tokens encoding the label, to understand if the attribution changed throughout the label tokens. The influence profiles (Fig. 5) reveal that the causal contribution localised to the embedded TATA motif (blue bands) is minimal — essentially indistinguishable from the surrounding non-discriminative background. The dominant peaks in the profile do not coincide with the biological signal; they recur periodically at the structural boundaries of the iCL prompt (the transitions into the label-run regions), tracking the syntactic [sequence] *→* [label_run] template rather than its semantic content.

Together, these analyses support the evidence that in-context learning in genomic models relies heavily on global positional structure rather than local motif extraction, with larger models prone to premature representational convergence across intermediate layers.

## 2. Discussion

In this work, we evaluated the in-context learning (iCL) capabilities of the Evo 2 suite across real-world and synthetic biological classification tasks. Our findings demonstrate that Evo 2 possesses genuine but strictly bounded iCL capabilities. The model achieves strong few-shot performance on short natural sequences — with Evo 2 7B reaching an F1 score of 0.902 on miRNA classification — outperforming parameter-matched, instruction-tuned medical LLMs. This confirms that nucleotide-level pretraining imparts domain-relevant representations that can be exploited at inference time without weight updates. However, three major boundaries constrain this capacity: sequence length, model scale, and prompt template dominance.

First, sequence length induces a qualitative failure mode in iCL that operates independently of biological signal complexity. On synthetic TATA-box tasks with identical classification signals, performance collapses from F1 = 0.629 at 20-100 bp to near chance (*F*1 *≤* 0.515) at 1-3 kb. Mechanistically, our Jacobian scope analysis reveals that at the kilobase scale, the model learns the syntax of the prompt without processing its sequence semantics. Furthermore, sequence length creates an inherent compute constraint: longer sequences consume significantly more memory per shot, truncating reachable example budgets (*N*) before asymptotic performance ceilings can be identified, as observed in AMR gene classification (~500-3,000 bp) and implementation limits during toxin evaluation. That said, we are careful to frame the entanglement with computational constraints. Longer sequences admit fewer in-context examples for the same compute — so that it marks the limit of what is currently feasible rather than a hard ceiling that more compute could not move.

Second, parameter scaling does not yield monotonic performance gains. Across all tasks, the 7B variant consistently outperformed the 40B model, a pattern preserved in the 20B spliced variant and pretrained base checkpoints. Our logit lens analysis provides a mechanistic rationale: while predictions in Evo 2 7B emerge gradually within the final 2-3 layers, Evo 2 40B exhibits premature representational convergence, maintaining saturated top-1 predictions across the entire second half of the network. This early commitment to next-token prediction appears to restrict the capacity of intermediate layers to maintain abstract, class-generalising representations necessary for few-shot discrimination. Testing this hypothesis via sparse autoencoder (SAE) probing or targeted layer interventions offers a clear avenue for future model architecture design.

Third, our results establish that raw sequence perplexity is an unreliable — and frequently misleading — proxy for downstream in-context classification accuracy. While perplexity decreases monotonically with *N* across almost all tasks, this drop reflects formatting and background adaptation rather than task discrimination. In the most severe case (TATA 1-3 kb), perplexity increased while classification F1 simultaneously collapsed. Evaluating genomic foundation models solely through perplexity risks drawing incorrect conclusions regarding downstream utility: future benchmarks should prioritise direct evaluation on classification tasks instead of next-token predictability.

As prompt engineering strategies refine and context efficiency improves, screening pipelines must adopt direct classification and adversarial robustness metrics rather than relying on single-pass perplexity filters. A clear example supporting this direction is found in recent literature, where researchers demonstrated how Evo 2 could be jailbroken to produce harmful sequences [19]. It’s important to note that those findings relied on a complex, multi-component apparatus that leveraged Evo 2 as well as other general and biological LMs. The positive iCL results open up a different solution, harder to detect and mitigate, and relying solely on Evo 2. The ability to guide a genomic model’s output class through incontext demonstrations could be exploited to generate adversarially ‘‘self-tuned’’ sequences, in a manner analogous to a GAN-based generation — that evade classifiers or bias model outputs towards desired functional categories.

## 3. Conclusion

Genomic foundation models open an exciting frontier for zero-shot and in-context biological reasoning, yet our findings reveal a fundamental gap between surface syntax and true biological semantics. On short sequences, in-context learning is a genuinely promising route: smaller models like Evo 2 7B rapidly pick up local patterns, and nucleotide-level pretraining instils representations that are immediately exploitable for motif discovery and short-sequence classification. At longer, multi-kilobase scales, this capacity breaks down. As hinted by the mechanistic analysis, models’ attention is absorbed by prompt formatting, rather than by the complex biological signal. Crucially, this failure is largely masked by perplexity: one of the field’s default evaluation metrics improves even as classification collapses, and so cannot be trusted as a proxy for downstream performance or genuine biological comprehension.

Nevertheless, our results show that models such as Evo 2 7B still solve many of the proposed tasks, despite these mechanistic behaviours. These architectures are at a very early stage and are being applied to a problem far harder than natural-language modelling, one in which the mapping from sequence to meaning is poorly characterised and even evaluating whether a model has truly learned remains unresolved. Part of the limitation may therefore be methodological rather than intrinsic: current frameworks for characterising biological sequences reflect prior, human-derived assumptions, and models that read structure directly from raw nucleotides may ultimately extend, rather than recapitulate, our understanding of the genome. Seen this way, today’s limitations are better understood as open problems than as fundamental barriers.

Two practical commitments follow. First, the short-sequence strengths that are useful now — in motif discovery and small-element engineering —warrant careful monitoring for dual-use screening of functional elements, extending the concerns recently raised by the scientific community [20] about oversight of AI-generated genomic designs following the publication of the first functional bacteriophage genome designed using Evo 2 [21]. Second, genomic foundation models should be evaluated directly on the downstream classification tasks they are meant to support, not on next-token predictability alone. To build models that reason across the full hierarchy of biology — from short regulatory motifs to multi-kilobase genomic architectures — the field must move past linear, text-based paradigms toward prompt strategies and architectures designed for the multi-dimensional language of life.

## 4. Methods

### 4.1. Models

The Evo 2 suite [3] comprises two families of models that differ in the amount of training they undergo. *Base* (pretrained) models were trained on sequences of up to 8k bases and have context windows — the maximum input length the model can process — ranging from 8k to 262k bases. They come in three sizes: 1, 7, and 40 billion parameters. *1M* (midtrained) models extend the base models with additional DNA sequences, reaching a context window of up to 1 million bases. The 1M models were originally released at 7B and 40B; a 20B version was later obtained by splicing the 40B model [3, 8], yielding a smaller model with near-identical capabilities (Supplementary Information S.1).

Because the base models offer only a limited context window, they restrict the number of in-context examples that can be supplied, especially for long sequences. We therefore used the 7B and 40B 1M models for all in-context learning experiments, and reserved the 1B base model exclusively for mechanistic interpretability comparisons. A direct comparison of the base and 1M variants on a length-unconstrained task is reported in Supplementary Information S.2.

Following previous work [11], we also included large language models as baselines. We focused on the MedGemma suite [22, 23] — specifically the 1.5 4B and 27B-text-only variants, both in their instruction-tuning (IT) version. This selection was based on two main criteria. First, the instruction-like nature of the task. Second, its medical nature and the specific training data for the models, which contained some high-level genomic knowledge. While MedGemma models are not genomic foundation models, as they lack nucleotide-level pretraining, they have extensive medical knowledge, including high-level genomic information, and their inclusion serves a dual control: matching parameter scale across small and large model tiers, and testing whether high-level biomedical domain knowledge is sufficient for in-context sequence classification.

### 4.2. Perplexity

In language modelling, perplexity (PPL) quantifies the uncertainty of a model when predicting a sequence of tokens *X* = (*x*_1_, *x*_2_,…, *x*_*T*_). Mathematically, it is defined as the exponentiated cross-entropy loss over the sequence:

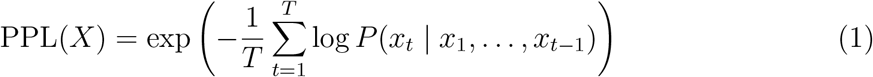

Intuitively, a perplexity of *K* means that at each token position, the model is as uncertain as if it were choosing uniformly at random among *K* equally likely options. In the context of In-Context Learning (ICL), evaluating raw sequence perplexity tests how well the model models the overall sequence distribution (e.g., nucleotide patterns, background sequence statistics, and prompt formatting), whereas downstream classification performance (such as macro F1 evaluated via perplexity scoring over discrete label completions) measures the model’s ability to discriminate between specific target classes.

### 4.3. In-context learning protocol

We adopted the approach proposed by Breslow and colleagues [11] for genomic sequences (see the paper’s Appendix E). For each test item, *N* examples per class were sampled without replacement from the training pool. Sequences were formatted as `[genomic_sequence] [buffer_sequence] [label_sequence]`, where the buffer and label sequences consisted of 24 repetitions of a single-nucleotide marker (e.g., GGGG…*×*24 for class 0, AAAA…*×*24 for class 1, CCCC…*×*24 for each sequence’s buffer). The query was formatted as `[genomic_sequence] [buffer_sequence] [candidate_label _sequence]`. Two forward passes were performed per test item — one for each candidate class label — and perplexity was computed over the label-run region only. Classification was assigned to the lower-perplexity candidate, and Macro F1 was adopted as the evaluation metric.

### 4.4. Datasets

We adopt three types of natural sequences: miRNA, Toxin-encoding genes, and AMR. miRNAs are small, naturally occurring non-coding RNAs involved in post-transcriptional gene regulation, acting as therapeutic targets and diagnostic or prognostic biomarkers (e.g., in cancer and cardiovascular disease) [24, 25]. Their short, well-characterised sequence and structural features make them an ideal benchmark to test whether iCL captures consistent, biologically grounded patterns from limited examples. Toxin-encoding genes are a class of direct biosecurity relevance: sequence-based recognition of toxin genes underlies existing biosecurity screening frameworks for DNA synthesis and gene ordering [26], and evaluating a gLM’s capacity to recognise these sequences shows both its potential misuse risks and its usefulness for bio-surveillance [27, 28]. Finally, AMR genes constitute a functionally defined but sequence-diverse class, frequently subject to horizontal gene transfer and rapid evolutionary turnover [29]; their accurate identification is central to genomic surveillance efforts addressing antimicrobial resistance, one of the top global public health threats [30].

Alongside the natural sequence dataset, we built two TATA-box datasets, distinguished only by the length of the generated DNA sequences. TATA-box sequences are well-characterised core promoter elements defined by a strongly conserved consensus motif [31]. Because they follow an explicit, well-established positional and compositional grammar, this task serves as a controlled benchmark. This allows us to isolate whether iCL can learn a known, well-defined sequence motif under near-ideal conditions, providing an interpretability anchor against which the more biologically complex, real-world classification tasks can be compared.

Each dataset was partitioned into an 80-20% train-test split. The training set served exclusively as the pool from which the contextual examples were sampled, following the in-context learning procedure described in Section 4.3, while the held-out test set was used for evaluation. This separation was adopted to guarantee a large sampling pool for the contextual examples and to prevent the reported results from being driven, either positively or negatively, by a specific outlier or a particular set of contextual examples.

#### miRNA

The dataset consists of 27,596 mature miRNA sequences from miRBase [32] (positive class), paired with 27,596 fragments of coding sequences from ESKAPEE pathogen reference genomes in NCBI RefSeq (negative class), length-matched to miRNA statistics (average ~26 bp). Negative sequences were selected to share similar GC background and length distribution while maintaining low sequence similarity to miRNAs (verified by pairwise alignment filtering). MicroRNAs (miRNAs) are short (~222-nucleotide) non-coding RNAs that regulate gene expression post-transcriptionally, primarily by guiding the RNA-induced silencing complex (RISC) to complementary sites in target mRNAs, leading to translational repression or degradation[33]. In the canonical biogenesis pathway, miRNA genes are transcribed from DNA into primary miRNA transcripts (pri-miRNAs), which are processed into precursor miRNAs (pre-miRNAs) and finally into mature miRNAs. Mature miRNAs typically interact with the 3’ untranslated region of target mRNAs [34].

#### Toxins

The dataset consists of 2,090 sequences (positive class) coding for either type I bacterial toxins or conotoxins, paired with 1,729 sequences coding for non-toxin proteins from ESKAPEE pathogen reference genomes in NCBI RefSeq (negative class), using a mixed-length distribution. Negative sequences were selected to share similar GC background and length distribution while maintaining low sequence similarity to toxins (verified by pairwise alignment filtering) Type I bacterial toxin non-redundant coding sequences were retrieved from the T1TAdb database [35] (length range: 27-300 nucleotides). Type I toxins are typically small, hydrophobic proteins that damage the bacterial cell membrane, restrict growth, or induce cell death. This helps bacteria adapt to stress, survive antibiotic treatment, or maintain plasmids. They are part of the type I toxin-antitoxin systems, i.e., genetic modules composed of a stable toxic protein and an unstable RNA antitoxin. The antitoxin acts as an antisense RNA to block the toxin’s translation. When the bacterium undergoes environmental stress, the antitoxin degrades, allowing the toxin to be produced [36]. Non-redundant conotoxin complete coding sequences were retrieved from ConoServer [37] (length range: 100-699 nucleotides). Conotoxins are a family of neurotoxic peptides naturally produced by venomous marine cone snails (genus *Conus*). These molecules primarily function by targeting and blocking specific ion channels and receptors in the nervous system. They evolved to immobilise and capture prey rapidly, but their extreme precision has made them promising tools in neuroscience and medicine [38].

#### AMR

The dataset consists of 8,302 antimicrobial resistance genes (ARGs) from the Pathogen Detection Reference Gene Catalogue [39] (positive class), paired with 8,302 non-ARG coding sequences from ESKAPEE reference genomes retrieved from the NCBI RefSeq [40] (negative class). Negative sequences were selected to share a similar genomic background and length distribution while maintaining low sequence similarity to ARGs (verified by pairwise alignment filtering). ARGs allow microorganisms, such as bacteria, to survive exposure to drugs designed to kill or inhibit them. Antimicrobial resistance (AMR) poses a major threat to public health worldwide, driven in part by the exchange of resistant bacteria and resistance genes across human, animal, and environmental reservoirs [41]. ESKAPEE is an acronym for a group of seven highly virulent and multidrug-resistant bacterial pathogens responsible for the majority of healthcare-associated infections worldwide. These bacteria are notorious for their ability to evade or ‘‘escape’’ conventional antibiotics [42]. The acronym stands for the following organisms, with NCBI RefSeq assembly IDs given in parentheses: *Enterococcus faecium* ((GCF_009734005.1), *Staphylococcus aureus* (GCF_000013425.1), *Klebsiella pneumoniae* (GCF_000240185.1), *Acinetobacter baumannii* (GCF_009035845.1), *Pseudomonas aeruginosa* (GCF_000006765.1), *Enterobacter cloacae* (as representative of *Enterobacter* spp., GCF_905331265.2), *Escherichia coli* (GCF_000005845.2)).

#### Synthetic TATA-box

The TATA-box is a highly conserved DNA sequence found in the promoter region of genes in eukaryotes and archaea. Named after its consensus sequence (typically with *TATA* motif). The canonical consensus sequence motif TATAWAWR (W = A/T, R = A/G) is located approximately 25 to 35 base pairs upstream of a gene’s transcription start site [43]. This synthetic dataset consists of 2,500 sequences of lengths drawn from *N* (*µ, σ*) with *µ* = (min + max)*/*2 and *σ* = (max *−* min)*/*6, ensuring ~99% of sequences lie within [min, max], paired with 2,500 sequence in which the consensus motif is absent. Two regimes: TATA-short (20-100 bp) and TATA-long (1-3 kb). Positive sequences contain a single TATAWAWR motif (W = A/T, R = A/G) embedded at a uniformly random position. Negative sequences have all TATAW occurrences destroyed by resampling the W position to C or G. Bases outside the motif are drawn uniformly from *{*A, T, C, G*}*.

### 4.5. Mechanistic interpretability

#### Logit lens

The logit lens [17] exploits the fact that a transformer’s output distribution is a fixed linear read-out of its final residual stream: applying that same unembedding read-out to the residual stream at an *intermediate* layer yields an interpretable, if partial, view of the model’s provisional next-token prediction at that depth. Because the architecture is a residual network, representations remain in a consistent basis across layers and are refined incrementally, which is what makes this cross-layer projection meaningful. In practice, at each layer we projected the (layer-normalised) residual-stream embedding at every token position to the vocabulary using the model’s unembedding matrix and recorded the top-1 token probability *P* (top-1), averaging across positions (token-wise average) to obtain a layer-by-model summary of the depth at which the network becomes confident in a single prediction. Rising top-1 probability across depth thus indexes how early the model commits to a concrete next-token guess rather than maintaining a more distributed, abstract representation.

#### Jacobian influence (Semantic Scope)

To quantify how strongly each input token causally shapes the model’s label prediction, we adopt the *Semantic Scope* variant of the Jacobian Scopes framework [18]. For a target label-run token, let *J*_*t*_ = *∂y/∂x*_*t*_ denote the Jacobian of the model’s final hidden state *y* with respect to the input embedding at position *t*. The Semantic Scope projects this Jacobian onto the unembedding direction *w* of the target token and takes its norm, yielding a per-token influence score *||w*^⊤^ *J*_*t*_||_2_; intuitively, this is the first-order change in the target logit induced by an infinitesimal perturbation of the token at position *t*, and thus measures the causal contribution of each context token to generating that target token. Unlike attention weights, which reflect where computation is routed rather than what causally drives the output, the Semantic Scope gives an exact, architecture-agnostic first-order attribution and requires only a single backward pass per target token. Influence profiles were computed over the last 24 label-run target tokens and summarised as averaged position-wise profiles for qualitative comparison across models. We note that, as a local first-order linearisation, the Semantic Scope captures direct token-level influence and does not resolve higher-order or layerwise interactions.

### 4.6. Computational resources

All experiments were run on a server with 8 H100 GPUs. The analyses were performed in Python, using common data science tools (Scikit-Learn, NumPy, PyTorch). Plots were generated in Python and refined iteratively with the help of large language models (see subsection “Use of large language models”)

### 4.7. Use of large language models

Large language models were used as assistive tools during this work. Specifically, they supported the generation and debugging of code for the experiments and analyses, and assisted in drafting the initial version of the manuscript and figure improvements. All model-generated content — code and text alike — was subsequently reviewed, edited, and verified by the authors.

## Supporting information

BA_etal_26_supplementary

## Data availability

Datasets for miRNA and AMR are derived from publicly available repositories (miRBase, NCBI). The synthetic TATA-box generation script is available upon request. Processed benchmark datasets will be deposited in a public repository upon acceptance.

## Code availability

The code to perform the analyses displayed in this paper is available at https://github.com/jrcf7/Evo2_iCL.

## Acknowledgements

The views expressed are those of the authors and do not necessarily reflect the official position of the European Commission. We would like to thank the colleagues of the Digital Health Unit (JRC.F7) at the Joint Research Centre of the European Commission for their helpful guidance and support. We thank Alessandro Zani, Lea Schuh, and Maddalena Querci for their comments and feedback on an early draft of the manuscript.

## Author contributions

Conceptualisation: L.B.A., F.C., M.P., G.L., M.R., V.R.S, S.C., M.C. Methodology: L.B.A., F.C. Computational investigation: L.B.A. Experimental investigation: L.B.A. Data curation: M.P., L.B.A. Visualisation: L.B.A., F.C., M.P. Supervision: S.C., M.C. Software: L.B.A. Formal analysis: L.B.A., F.C. Writing — original draft: L.B.A., M.P. Writing — review and editing of the final version of the paper: L.B.A., F.C., M.P., G.L., M.R., V.R.S, S.C., M.C.

## References

[1] Consens, M. E., Dufault, C., Wainberg, M., Forster, D., Karimzadeh, M., Goodarzi, H., Theis, F. J., Moses, A., & Wang, B. Transformers and genome language models. Nature Machine Intelligence, 7(3), 346–362, 2025. 10.1038/s42256-025-01007-9.

[2] Guo, F., Guan, R., Li, Y., Liu, Q., Wang, X., Yang, C., & Wang, J. Foundation models in bioinformatics. National Science Review, 12(4), nwaf028, 2025. 10.1093/nsr/nwaf028.

[3] Brixi, G., Durrant, M. G., Ku, J., Naghipourfar, M., Poli, M., Sun, G., Brockman, G., et al. Genome modelling and design across all domains of life with Evo 2. Nature, pp. 1–13, Mar. 2026. 10.1038/s41586-026-10176-5.

[4] Veiner, M. & Supek, F. The DNA dialect: a comprehensive guide to pretrained genomic language models. Molecular Systems Biology, 22(3), 309–332, 2026. 10.1038/s44320-025-00184-4.

[5] Nguyen, E., Poli, M., Faizi, M., Thomas, A., Birch-Sykes, C., Wornow, M., Patel, A., et al. HyenaDNA: long-range genomic sequence modeling at single nucleotide resolution. arXiv, 2023. 10.48550/arXiv.2306.15794.

[6] Nguyen, E., Poli, M., Durrant, M. G., Kang, B., Katrekar, D., Li, D. B., Bartie, L. J., et al. Sequence modelling and design from molecular to genome scale with Evo. Science, 386(6723), eado9336, 2024. 10.1126/science.ado9336.

[7] Sanabria, M., Hirsch, J., Joubert, P. M., & Poetsch, A. R. DNA language model learns sequence context in the human genome. Nature Machine Intelligence, 6(8), 911–923, 2024. 10.1038/s42256-024-00872-0.

[8] Arc Institute. Evo2 v0.5.0: Evo 2 20B release, 2026. https://github.com/ArcInstitute/evo2/releases/tag/v0.5.0, xGitHub, ArcInstitute/evo2.

[9] McInerney, J. O. Genomic perplexity and the evolution of context-dependent function. Molecular Biology and Evolution, 43(3), msag041, 03 2026. 10.1093/molbev/msag041.

[10] Wei, B., Che, Z., Li, N., Götting, J., Nedungadi, S., Michael, J., Yue, S., Hendrycks, D., Henderson, P., Wang, Z., Donoughe, S., & Mazeika, M. Best Practices for Biorisk Evaluations on Open-Weight Bio-Foundation Models. in NeurIPS 2025 Workshop on Biosecurity Safeguards for Generative AI, 2025. https://openreview.net/forum?id=N5DhtpYJ21.

[11] Breslow, N., Mishra, A., Schatz, M., Liu, A., Revsine, M., & Khashabi, D. Genomic Next-Token Predictors are In-Context Learners. Transactions on Machine Learning Research, 2026. https://openreview.net/forum?id=KmNFx8DmaZ.

[12] Zhang, P., Prabhu, S., Grama, G., Bang, S., & Lee, H. Self-Contemplating In-Context Learning Enhances T Cell Receptor Generation for Novel Epitopes. in Knowles, D. A. & Koo, P. K. (eds), Proceedings of the 20th Machine Learning in Computational Biology meeting, Vol. 311, pp. 251--269, New York, USA, Sep. 2025. Proceedings of Machine Learning Research. https://proceedings.mlr.press/v311/zhang25a.html.

[13] Li, Q., Zhan, Z., Feng, S., Zhu, Y., He, Y., Wu, W., Shi, Z., et al. Functional in-context learning in genomic language models with nucleotide-level supervision and genome compression. bioRxiv, 2026. 10.64898/2026.01.27.702015.

[14] Kapoor, A. & Lee, W. In-Context Learning in Genomic Language Models as a Biological Evaluation Task, Dec. 2025. 10.21203/rs.3.rs-8287395/v1, Research Square Preprint.

[15] Brown, T. B., Mann, B., Ryder, N., Subbiah, M., Kaplan, J., Dhariwal, P., et al. Language models are few-shot learners. in Advances in Neural Information Processing Systems, Vol. 2020-December, 2020.

[16] Wang, L., Chen, X., Deng, X., Wen, H., You, M., Liu, W., Li, Q., & Li, J. Prompt engineering in consistency and reliability with the evidence-based guideline for LLMs. npj Digital Medicine, 7(1), 2024. 10.1038/s41746-024-01029-4.

[17] nostalgebraist. Interpreting GPT: the logit lens, 2020. https://www.lesswrong.com/posts/AcKRB8wDpdaN6v6ru/interpreting-gpt-the-logit-lens, LessWrong/AI Alignment Forum.

[18] Liu, T. J. B., Zadeoğlu, B., Boullé, N., Sarfati, R., & Earls, C. J. Jacobian Scopes: token-level causal attributions in LLMs, 2026. https://arxiv.org/abs/2601.16407.

[19] Zhang, Z., Zhou, Z., Jin, R., Cong, L., & Wang, M. GeneBreaker: Jailbreak Attacks against DNA Language Models with Pathogenicity Guidance. in The Fourteenth International Conference on Learning Representations, 2026. https://openreview.net/forum?id=C5OIolrNJd.

[20] Inglesby, T. V. & Hanke, M. S. AI-designed viral genomes. Science, 393(6811), 563–564, 2026. 10.1126/science.aej8512.

[21] King, S. H. et al. Generative design of bacteriophages with genome language models. Science, 393(811), eaec2657, 2026. 10.1126/science.aec2657.

[22] Sellergren, A., Kazemzadeh, S., Jaroensri, T., Kiraly, A., Traverse, M., Kohlberger, T., Xu, S., Jamil, F., Hughes, C., Lau, C., Chen, J., Mahvar, F., Yatziv, L., Chen, T., Sterling, B., Baby, S. A., Baby, S. M., Lai, J., Schmidgall, S., Yang, L., Chen, K., Bjornsson, P., Reddy, S., Brush, R., Philbrick, K., Asiedu, M., Mezerreg, I., Hu, H., Yang, H., Tiwari, R., Jansen, S., Singh, P., Liu, Y., Azizi, S., Kamath, A., Ferret, J., Pathak, S., Vieillard, N., Merhej, R., Perrin, S., Matejovicova, T., Ramé, A., Riviere, M., Rouillard, L., Mesnard, T., Cideron, G., bastien Grill, J., Ramos, S., Yvinec, E., Casbon, M., Buchatskaya, E., Alayrac, J.-B., Lepikhin, D., Feinberg, V., Borgeaud, S., Andreev, A., Hardin, C., Dadashi, R., Hussenot, L., Joulin, A., Bachem, O., Matias, Y., Chou, K., Hassidim, A., Goel, K., Farabet, C., Barral, J., Warkentin, T., Shlens, J., Fleet, D., Cotruta, V., Sanseviero, O., Martins, G., Kirk, P., Rao, A., Shetty, S., Steiner, D. F., Kirmizibayrak, C., Pilgrim, R., Golden, D., & Yang, L. MedGemma Technical Report, 2026. https://arxiv.org/abs/2507.05201.

[23] Sellergren, A., Gao, C., Mahvar, F., Kohlberger, T., Jamil, F., Traverse, M., Tono, A., Sadjad, B., Yang, L., Lau, C., Yatziv, L., Chen, T., Sterling, B., Philbrick, K., Tiwari, R., Liu, Y., Jajoo, M., Sankarapu, C., Vispute, S., Purandare, H., Mishra, A. B., Schmidgall, S., Tu, T., Palepu, A., Park, C., Strother, T., Thapa, R., Cheng, Y., Singh, P., Black, K., Matias, Y., Chou, K., Hassidim, A., Goel, K., Barral, J., Warkentin, T., Shetty, S., Webster, D., Virmani, S., Steiner, D. F., Kirmizibayrak, C., & Golden, D. MedGemma 1.5 Technical Report, 2026. https://arxiv.org/abs/2604.05081.

[24] Laggerbauer, B. & Engelhardt, S. MicroRNAs as Therapeutic Targets in Cardiovascular Disease. Journal of Clinical Investigation, 2022. 10.1172/JCI159179.

[25] Jurj, A., Dragomir, M. P., Li, Z., & Calin, G. A. MicroRNAs in Oncology: A Translational Perspective in the Era of AI. Nature Reviews Clinical Oncology, 23(4), 239–259, 2026. 10.1038/s41571-025-01114-x.

[26] Wheeler, N., Carter, S. R., Alexanian, T., Isaac, C., Millett, P., & Yassif, J. Overcoming Challenges to Developing a Common Global Baseline for Nucleic Acid Synthesis Screening. Applied Biosafety, 2024. PMCID: PMC11313551.

[27] Bloomfield, D., Pannu, J., Zhu, A. W., Ng, M. Y., Lewis, A., Bendavid, E., et al. AI and Biosecurity: The Need for Governance. Science, 385(6711), 831–833, 2024. 10.1126/science.adq1977.

[28] Bloomfield, D., Pannu, J., Zhu, A. W., Ng, M. Y., et al. Dual-Use Capabilities of Concern of Biological AI Models. PLOS Computational Biology, 2025. PMCID: PMC12061118.

[29] Darby, E. M., Trampari, E., Siasat, P., Gaya, M. S., Alav, I., Webber, M. A., & Blair, J. M. Molecular mechanisms of antibiotic resistance revisited. Nature Reviews Microbiology, 21(5), 280–295, 2023.

[30] World Health Organization. Antimicrobial Resistance, 2023. https://www.who.int/news-room/fact-sheets/detail/antimicrobial-resistance, Accessed: 2026-08-05.

[31] Smale, S. T. & Kadonaga, J. T. The RNA Polymerase II Core Promoter. Annual Review of Biochemistry, 72, 449–479, 2003. 10.1146/annurev.biochem.72.121801.161520.

[32] Kozomara, A., Birgaoanu, M., & Griffiths-Jones, S. miRBase: from microRNA sequences to function. Nucleic Acids Research, 47(D1), D155–D162, 2019. 10.1093/nar/gky1141.

[33] Bartel, D. P. Metazoan MicroRNAs. Cell, 173(1), 20–51, mar 2018. 10.1016/j.cell.2018.03.006.

[34] Shang, R., Lee, S., Senavirathne, G., & Lai, E. C. microRNAs in action: biogenesis, function and regulation. Nature Reviews Genetics, 24(12), 816–833, 2023. 10.1038/s41576-023-00611-y.

[35] Tourasse, N. J. & Darfeuille, F. T1TAdb: the database of type I toxin--antitoxin systems. RNA, 27(12), 1471–1481, 2021. 10.1261/rna.078802.121.

[36] Nonin-Lecomte, S., Fermon, L., Felden, B., & Pinel-Marie, M.-L. Bacterial Type I Toxins: Folding and Membrane Interactions. Toxins, 13(7), 490, 2021. 10.3390/toxins13070490.

[37] Kaas, Q., Westermann, J.-C., Halai, R., Wang, C. K. L., & Craik, D. J. ConoServer, a database for conopeptide sequences and structures. Bioinformatics, 24(3), 445–446, 2008. 10.1093/bioinformatics/btm596.

[38] Li, R., Yu, J., Ye, D., Liu, S., Zhang, H., Lin, H., Feng, J., & Deng, K. Conotoxins: Classification, Prediction, and Future Directions in Bioinformatics. Toxins, 17(2), 78, 2025. 10.3390/toxins17020078.

[39] Feldgarden, M., Brover, V., Gonzalez-Escalona, N., Frye, J. G., Haendiges, J., Haft, D. H., Hoffmann, M., Pettengill, J. B., Prasad, A. B., Tillman, G. E., Tyson, G. H., & Klimke, W. AMRFinderPlus and the Reference Gene Catalog facilitate examination of the genomic links among antimicrobial resistance, stress response, and virulence. Scientific Reports, 11, 12728, 2021. 10.1038/s41598-021-91456-0.

[40] O’Leary, N. A., Wright, M. W., Brister, J. R., Ciufo, S., Haddad, D., McVeigh, R., Rajput, B., Robbertse, B., Smith-White, B., Ako-Adjei, D., & et al. Reference sequence (RefSeq) database at NCBI: current status, taxonomic expansion, and functional annotation. Nucleic Acids Research, 44(D1), D733–D745, 2016. 10.1093/nar/gkv1189.

[41] Larsson, D. G. J. & Flach, C.-F. Antibiotic resistance in the environment. Nature Reviews Microbiology, 20(5), 257–269, 2022. 10.1038/s41579-021-00649-x.

[42] Oyenuga, N., Cobo-Díaz, J. F., Alvarez-Ordóñez, A., & Alexa, E.-A. Overview of Antimicrobial Resistant ESKAPEE Pathogens in Food Sources and Their Implications from a One Health Perspective. Microorganisms, 12(10), 2084, 2024. 10.3390/microorganisms12102084.

[43] Bucher, P. Weight matrix descriptions of four eukaryotic RNA polymerase II promoter elements derived from 502 unrelated promoter sequences. Journal of Molecular Biology, 212(4), 563–578, 1990. 10.1016/0022-2836(90)90223-9.

