## Supplementary material for "Evo 2 as a classification machine: evidence from in-context learning and mechanistic interpretability": BA_etal_26_supplementary

### Supplementary Information

#### S.1. Evo 2 20B matches the 40B model on in-context learning

The 1M (midtrained) Evo 2 suite was originally released at 7B and 40B parameter scales; the intermediate 20B variant was subsequently obtained by splicing the 40B model rather than by independent training [3, 8]. Because the 20B model is derived directly from the 40B weights, it is expected to inherit the 40B model's behaviour, and we treated the two as a single design point in the main text (Table 2) to avoid redundancy. Here we report the head-to-head comparison that justifies this choice.

Figure S1 contrasts the shot-scaling behaviour of the 20B and 40B models across the evaluated tasks. The two curves are nearly indistinguishable: the 20B model tracks the 40B trajectory across the full range of $N$, converging to peak macro F1 within a few thousandths of the 40B result (e.g., miRNA $\mathrm{F1} = 0.826$ for 20B versus 0.846 for 40B) and reproducing the same plateau-and-degradation pattern beyond $N \approx 64$. Crucially, the 20B model shows none of the benefits one might hope for from a smaller-but-independent model — it does not recover the 7B model's continued improvement — confirming that it behaves as a compressed copy of the 40B model rather than as a distinct scale. This near-identity has a practical upshot: the 20B model offers the 40B model's iCL behaviour at reduced memory and compute cost, but confers no accuracy advantage, and neither variant closes the gap to the 7B model.

#### S.2. Base (pretrained) models underperform the 1M (midtrained) models

The main-text experiments use the 1M (midtrained) Evo 2 models, which extend the base (pretrained) checkpoints with additional long-sequence DNA training and reach a 1M-base context window (Methods). To assess whether the additional midtraining matters for in-context learning — as opposed to simply enabling longer contexts — we compared the base and 1M variants on the miRNA task, where the short sequence length keeps every model well within its context budget and thus permits a fair comparison across both training regimes and all parameter scales.

Figure S2 shows that the base models are substantially worse in-context learners than their

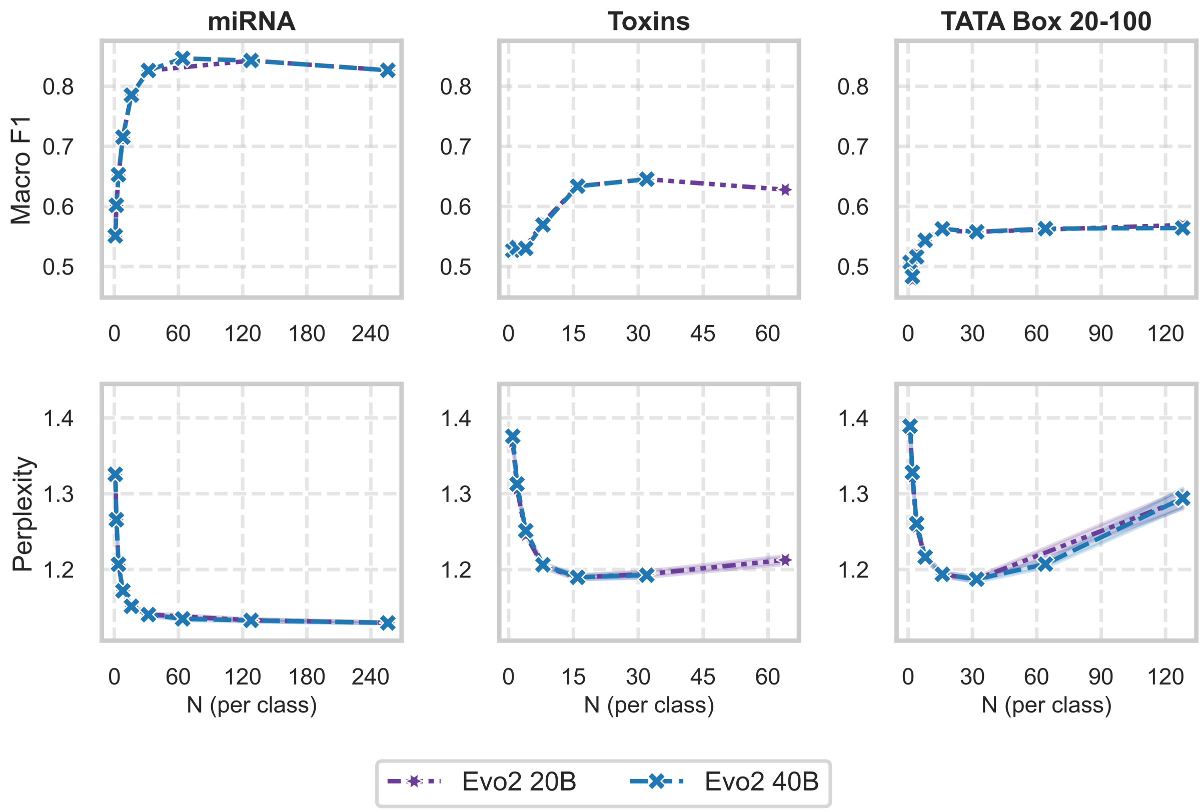


**Supplementary Figure S1. Evo 2 20B versus 40B across in-context learning tasks.** Macro F1 as a function of the number of in-context examples per class ($N$) for the spliced 20B model and the 40B model from which it is derived.

1M counterparts. At matched parameter scale, the 1M models consistently outperform the base models across the range of $N$, indicating that the long-sequence midtraining stage — and not merely the enlarged context window — confers representations that iCL can exploit. This is consistent with the main-text observation (Fig. 2) that the 1M-context miRNA variant outperforms its base counterpart.

Two ordering effects nonetheless persist across training regimes. First, the 7B model remains the strongest performer even among the base checkpoints: the base 7B model still surpasses the larger base models, mirroring the 7B > 40B ordering seen in the 1M suite and reinforcing that the 7B scale, rather than the largest scale, is best suited to this perplexity-scoring protocol. Second, the base-versus-1M gap does not close with scale, so the advantage of midtraining is not an artefact of any single model size. Together, these results show that both mid-training and the choice of the 7B scale contribute to iCL performance, and that neither factor can be recovered by the other.

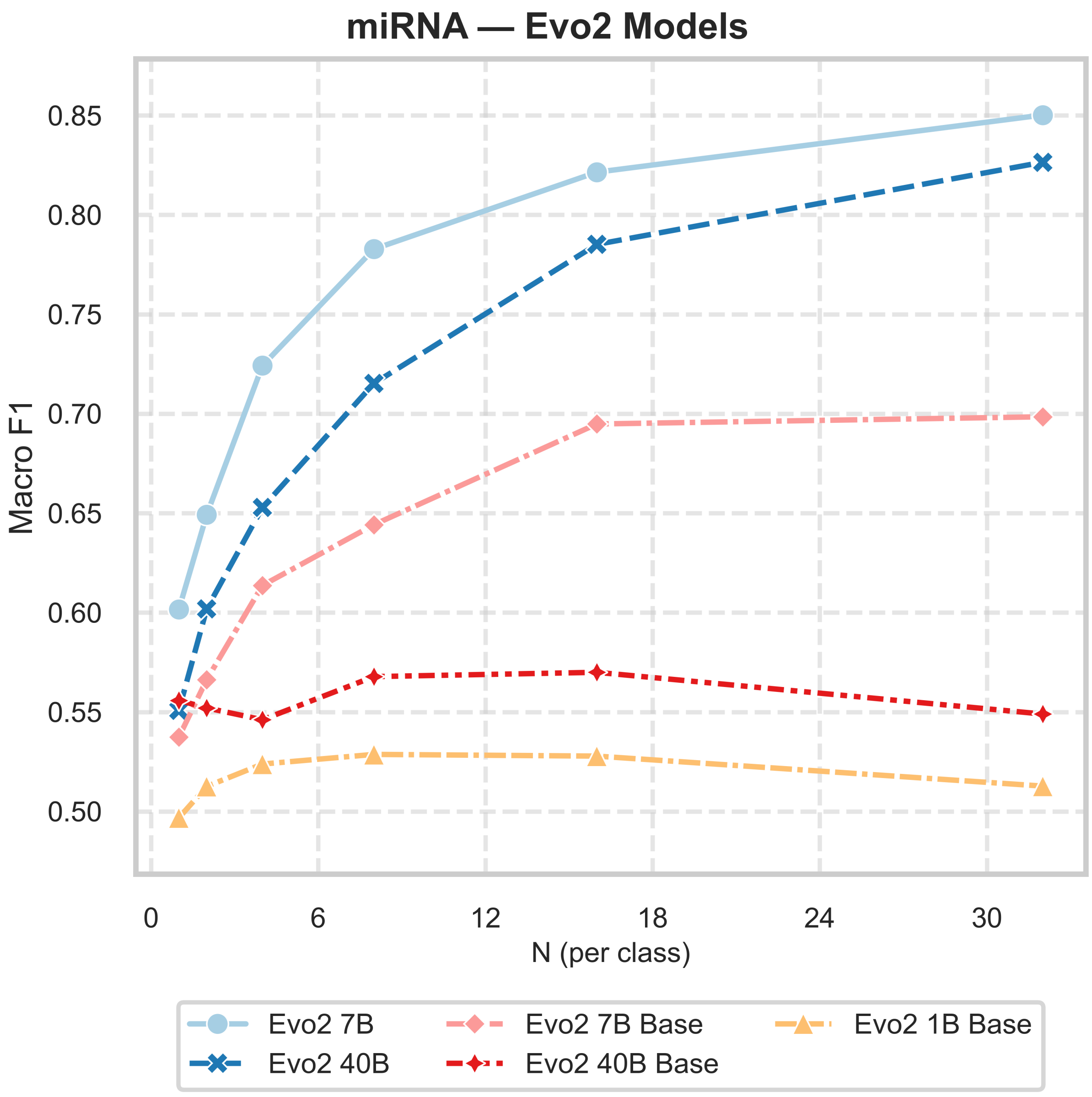


**Supplementary Figure S2. Base versus 1M Evo 2 models on miRNA in-context learning.** Macro F1 as a function of the number of in-context examples per class ($N$) for the base (pretrained) and 1M (midtrained) Evo 2 models, evaluated on the miRNA task. Coloured lines distinguish models and training regimes as indicated in the legend.
